# Microstructure Aware Modeling Of Biochemical Transport In Arterial Blood Clots

**DOI:** 10.1101/2021.01.25.428179

**Authors:** Chayut Teeraratkul, Debanjan Mukherjee

## Abstract

Flow-mediated transport of biochemical species is central to thrombotic phenomena. Comprehensive three-dimensional modeling of flow-mediated transport around realistic macroscale thrombi poses challenges owing to their arbitrary heterogeneous microstructure. Here, we develop a microstructure aware model for species transport within and around a macroscale thrombus by devising a custom preconditioned fictitious domain formulation for thrombus-hemodynamics interactions, and coupling it with a fictitious domain advection-diffusion formulation for transport. Microstructural heterogeneities are accounted through a hybrid discrete particle-continuum approach for the thrombus interior. We present systematic numerical investigations on unsteady arterial flow within and around a three-dimensional macroscale thrombus; demonstrate the formation of coherent flow structures around the thrombus which organize advective transport; illustrate the role of the permeation processes at the thrombus boundary and subsequent intra-thrombus transport; and characterize species transport from bulk flow to the thrombus boundary and vice versa.

## 1 Introduction

Pathological blood clotting or thrombosis is the primary underlying cause of most major cardiovascular diseases including ischemic stroke^[52]^. Despite recent medical advances, the W.H.O. estimates that nearly 5.7 million people worldwide are killed by stroke^[36,47]^; making it a leading global cause of death and disability. Blood flow and transport processes in the vicinity of a large-artery thrombus (*pathologically formed clot*) are central to thrombus growth^[34,16]^, mechanics^[4,9,17]^, and lysis^[15,6]^. Of particular interest is the near-thrombus transport of biochemical species that play key roles in thrombotic phenomena and thrombolysis. Specific examples include: (a) transport of agonists like adenosine diphosphate (ADP) and thromboxane (TxA2) released from thrombus site which can further activate platelets^[20]^ and enable disease progression; (b) delivery of thrombolytic drug tissue plasminogen activator (tPA) into an occlusive thrombus^[48,29]^; and (c) gas transport phenomena in intraluminal thombi formed in aneurysms^[53]^. Thus, comprehensive understanding of flow-mediated biochemical species transport in and around a macroscopic large-artery thrombus is critical for understanding disease scenarios and evaluating treatment efficacy.

Computational approaches offer a viable avenue to investigate such phenomena. Flow and transport processes in thrombi have been studied at the microscale using non-equilibrium particle based approaches^[39,56,57,43,46,58]^, as well as at the macroscale using continuous media models^[23,24,18,38,31,60]^. Several of these macroscale models have focused on biochemical transport from the perspective of coagulation agonist transport^[24,16]^, while thrombolysis has received lesser attention^[37,3,38]^. Extending such modeling approaches to threedimensional large artery thrombi continues to pose challenges, and comprehensive three-dimensional unsteady computational models are rare, especially since real human thrombi are known to be of heterogeneous micro-composition^[54,51,61,50]^. A microstructure aware approach for comprehensive characterization of flow-mediated biochemical transport in macroscale thrombi is of major significance. For example, in thrombolytic drug delivery, unlike conventional advection-diffusion-reaction processes in continuous media, thrombolysis progresses heterogeneously with observed dependency on microstructural features such as packing density, and fibrin network architecture^[2,10,49,6]^. In a series of prior works we have proposed a mesoscale hybrid particle-continuum approach for handling arbitrary thrombus shape and microstructure for macroscopic arterial thrombi^[33,44,45]^ to address this aforementioned challenge. We demonstrated the efficacy of this approach in handling thrombus-hemodynamics interactions for two-dimensional models of large-artery thombi. Here we employ this approach to devise a heterogeneous, microstructure aware model for unsteady multiscale biochemical species transport within and around a three-dimensional large-artery thrombus.

## 2 Methods

### 2.1 Modeling arterial flow around a thrombus

Unsteady flow around a heterogeneous thrombus aggregate is computed using a hybrid particle-continuum fictitious domain finite element approach outlined in prior work^[33,45]^. Briefly, we consider the total computational domain Ω = Ω_*f*_ ⊕ Ω_*t*_, with Ω_*f*_ as the fluid (*blood*) domain, and Ω_*t*_ the thrombus domain with heterogeneous microstructure composed of a union of mesoscopic discrete particle domains 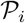 such that: 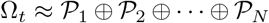. Blood is assumed to be an incompressible, Newtonian fluid with viscosity *μ_f_* and density *ρ_f_*. The Navier-Stokes momentum and mass balance equations are solved using a Petrov-Galerkin stabilized formulation^[33,45]^, with the overall weak form stated as follows:

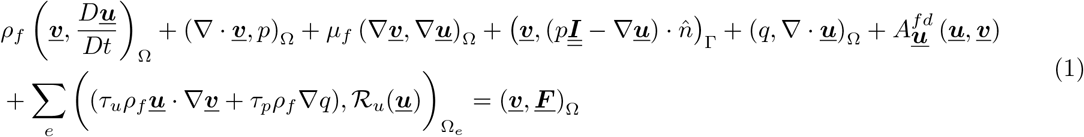

where we use the notation 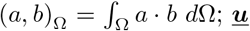, *p* are the velocity and pressure trial functions; ***<u>υ</u>***, *q* are the respective test functions; 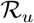 is the residual of the momentum and continuity equations; *τ_u_* and *τ_p_* are the velocity and pressure stabilization terms^[11]^; and ***<u>F</u>*** denotes the external body force (*gravity*). The term 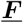 denotes the interaction contribution for the fictitious domain which constrains the velocity ***<u>u</u>*** to be ***<u>u</u>***_0_ and is defined as:

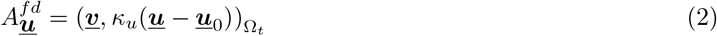

and *κ_u_* is a velocity penalization term over the fictitious domain modeled as^[33]^:

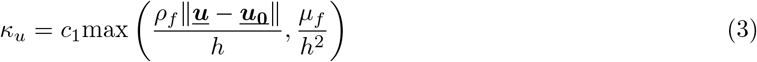

where *h* is the element size. Efficient resolution of the fictitious particle domain requires the mesh sizing in the fictitious domain to be commensurate with particle size^[44]^. With arbitrary three-dimensional microstructures, this sizing requirement can lead to a fine fictitious domain mesh, and a monolithic solution of associated velocity and pressure fields leads to a large linear system. The associated memory requirements make iterative Krylov solvers the strategy of choice, yet the asymmetric linear system resulting from stabilized formulation in Equation 1 can raise the required number of Krylov iterations^[40]^. Consequently, large three-dimensional problems with complex microstructures can become prohibitively expensive to solve. To mitigate this problem, here we devise a custom preconditioner based on the pressure convection diffusion (PCD) scheme^[35,12,14]^ to extend our fictitious domain method (*referred to as PCD-FD*). Briefly, following an implicit time discretization and linearization using Newton iteration, the matrix problem for Equation 1 can be written as:

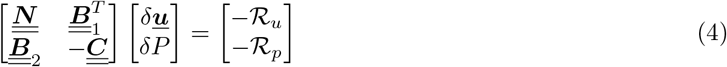

where *δ**<u>u</u>*** and *δP* are solution increments in the Newton iteration, and 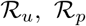 are the iteration residuals as stated before. PCD preconditioning seeks to approximate the pressure Schur complement arising from the following block LU decomposition of Equation 4:

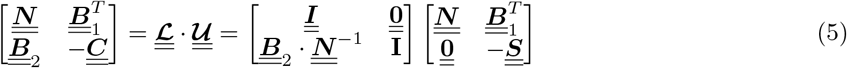

where the pressure Schur compliment is 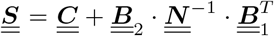. The upper-triangular matrix 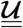 can be viewed as a preconditioner, and if 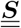 is computed exactly, then convergence with GMRES solver is guaranteed in 2 steps^[14]^. This is computationally expensive as it involves computing 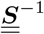, a dense matrix inversion. PCD preconditioning strategy approximates the action of 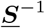 by the following:

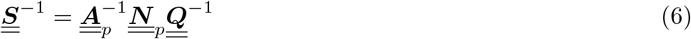

where 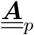 and 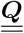 are the pressure Laplacian operator and velocity mass matrix respectively. The matrix 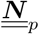 is the action of operator 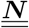 projected onto the pressure basis functions. The PCD preconditioner procedure thus contains a mass matrix solve, a matrix-vector multiplication, and a pressure Laplacian solve - which we implemented using a FEniCS^[28]^ based library FENaPack^[5]^.

### 2.2 Modeling species transport

In prior works^[33,44,45]^ our fictitious domain approach focused on resolving unsteady thrombus-hemodynamics interactions. Here, we extend this approach to couple unsteady flow with a custom fictitious domain advection-diffusion (AD) formulation for biochemical species transport. We devised a Petrov-Galerkin stabilized finite element formulation for the classical AD equation to stabilize for flow regimes with high Peclet number; along with an added contribution from fictitious domain interaction. The resultant weak formulation is stated as below using the same notation as in Equation 1:

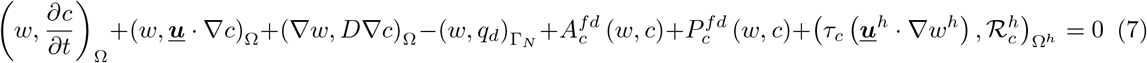

Here *c* is the concentration trial-function; *w* the test-function; *D* the diffusivity of the biochemical species of interest; ***<u>u</u>*** is the local velocity field (*obtained from the flow problem discussed in Section 2.1*); and 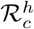 denotes the discretized AD equation residual. The last term in Equation 7 denotes Petrov-Galerkin (SUPG) stabilization, with *τ_c_* as the stabilization parameter. The term 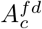 denotes the fictitious domain interaction contribution. Following the velocity formulation, here we set this contribution in form of a penalty term that constrains the species concentration (*c*) inside the fictitious domain to a set concentration *c*_0_, such that:

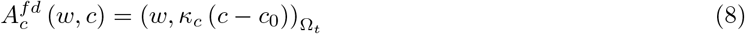

where *κ_c_* is the equivalent penalty parameter for the concentration problem; modeled as^[13]^:

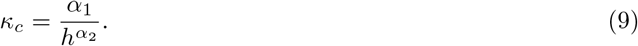

Furthermore, we note that particles composing Ω_*t*_ are not actual platelets/cells, but mesoscopic computational units used for discretized reconstruction of the heterogeneous thrombus domain^[33]^. While these particles are assumed to inherently hinder flow in the interior on account of them physically occupying the fictitious domain space, they are numerically allowed to contribute to the species concentration field. This contribution is accounted for in Equation 7 through a particle source term 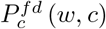, modeled as follows:

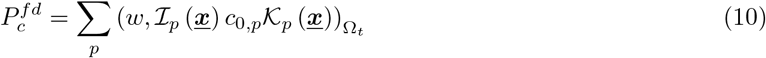

where *p* denotes index for the particle in 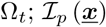 is an indicator function with value unity if 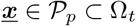and 0 otherwise; *c*_0,*p*_ is the strength of the source term; and 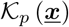 is a kernel function used to impose the source on the domain 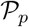. The kernel function is chosen to satisfy two mathematical conditions: (a) the kernel must have a support equivalent to the size of the particle 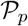; and (b) within the particle domain 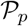 the kernel must integrate to 1.0. The kernel function is, thus, similar in notion to kernel based representations in Reproducing Kernel Particle Methods (RKPM)^[25,26]^, RKPM kernels in Immersed Finite Element Method (IFEM)^[59,27]^, and Smoothed Particle Hydrodynamics (SPH)^[32]^ in obtaining a meshfree particle based interpolation. Here we use a truncated Gaussian function for our kernel.

### 2.3 Quantifying coherent flow structures around thrombus

Coherent flow structures around the thrombus were quantified by computing the Finite Time Lyapunov Exponent (FTLE) field from simulated flow velocity data for the final cardiac cycle, as described in prior works^[41,45]^. Briefly, trajectories of a collection of massless tracers seeded on a Cartesian grid around the thrombus, were computed by numerically integrating the flow velocity interpolated at tracer locations as follows:

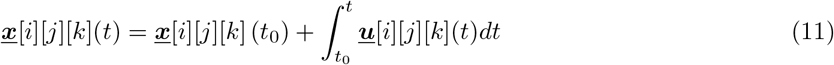

where [*i*][*j*][*k*] denotes indexing based on the Cartesian grid. Using the original seed coordinates 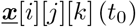 as the reference configuration, and the integrated tracer coordinates at time 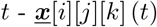 - as the current configuration, we computed a deformation gradient 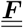 for tracer kinematics as follows^[41,45]^:

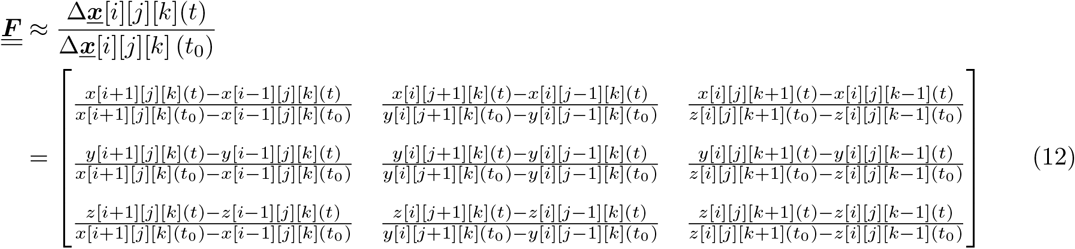

The FTLE field was then computed based on the scaled maximal eigenvalues of the Cauchy-Green deformation tensor 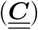 associated with the above deformation gradient 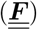 as follows:

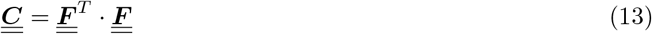

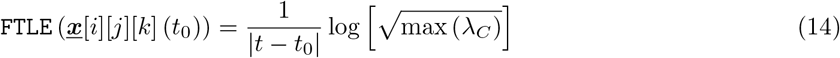

with λ_*C*_ denoting eigenvalues of the tensor 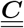. The computed FTLE field was mapped back onto the original Cartesian grid where tracers were seeded for visualization.

### 2.4 Numerical experiments on three-dimensional model system

We conducted a series of quantitative analyses on a simplified, yet representative, 3-dimensional model system. The system comprises a cylindrical vessel of diameter 6.0 *mm* (*equivalent of the human common carotid artery*) as shown in Figure 1. A physiologically realistic pulsatile flow profile obtained from prior studies^[22]^ was imposed at the inlet to drive flow through the vessel (*profile shown in Figure 2.b*.). A simplified model thrombus domain (Ω_*t*_) was created in form of a hemispherical shell of radius 3.0 *mm* booleaned with the vessel using OpenSCAD^[1]^ (see *Figure 1*). Corresponding discrete particle thrombus reconstruction was generated using tessellation based particle algorithms outlined in^[33,45]^. The resulting heterogeneous microstructure had an average discrete particle size of 98 microns (see *also supplementary material*), and average pore-fraction 0.68, with core porosity much lower than shell porosity. Detailed microstructural characterization, including pore-size and permeability estimates, was not conducted within the scope of this study. However, a simplified baseline permeability estimate for the discrete particle aggregate was obtained using classical Kozeny-Carman relation to be around 197 *μ*m^2^, which was reasonably within the regime of previously reported values^[55]^.

**Figure 1:**
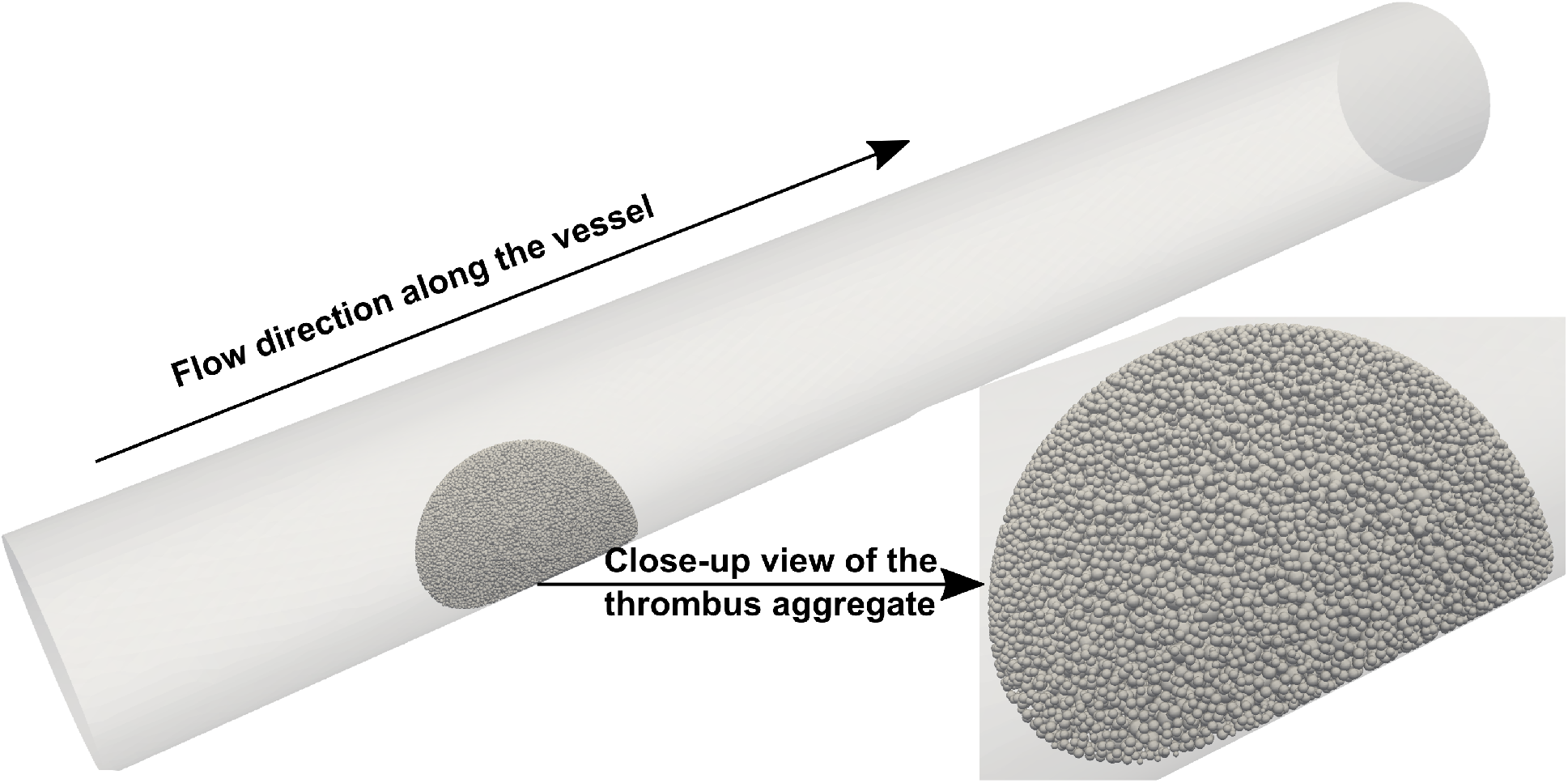
Illustration of the three-dimensional arterial thombus model used for the study, with a close-up view of the discrete particle representation of the thrombus aggregate.

Unstructured tetrahedral mesh with target edge size 100.0 *μm* was created for the vessel. A local mesh refinement region around Ω_*t*_ was created with a target edge size of 40.0 *μm*. Refinement mesh sizing was chosen such that histogram of element sizes had a lower mode than histogram of discrete particle sizes in Ω_*t*_^[44]^. Blood flow around the thrombus was simulated for three cardiac cycles using the PCD-FD method (see *Section 2.1*) with a cycle period of 0.90 *sec* and numerical time-step = 0.001 *sec*. Flow velocity field from the final cardiac cycle was used to compute FTLE fields (see *Section 2.3*) using a Cartesian grid of tracers of resolution 0.5 *mm*. The final cycle flow velocity field was further used to compute species concentration using the fictitious domain AD formulation (*see Section 2.2*). Species of varying diffusivity values starting from a physiologic range of 5 × 10^−5^*mm*^2^/*s*^[38]^ were studied. For each species we simulated two cases: (a) imposing a fixed species concentration at the vessel inlet; and (b) imposing fixed species concentration for discrete particles along the thrombus boundary within a thickness of 0.2 *mm* (≈ 6.67 % *of thrombus radius*). To ensure species transport simulations run over a sufficiently long duration, the advection-diffusion problem was simulated for 40 cardiac cycles by looping the flow velocity data from the final cycle. Finally, the cycle velocity data were averaged over time, to run steady-state versions of the fictitious domain AD formulation for long-term intra-thrombus transport behavior.

## 3 Results

### 3.1 Flow within and around the thrombus

Unsteady flow patterns were visualized for flow data from the final cardiac cycle as presented in Figure 2. For illustrating three-dimensional extra-thrombus flow patterns, streamlines at four instances during the cardiac cycle were generated (*Figure 2, panel b*.) - denoting mid-systole (*T*1), peak systole (*T*2), mid-deceleration (*T*3), and diastole (*T*4). The streamlines are presented in panel c., demonstrating the efficacy of our PCD-FD approach in resolving thrombus-hemodynamics interactions for three-dimensional heterogeneous thrombus aggregates. Furthermore, the PCD-FD formulation enables characterizing both extra-thrombus and intrathrombus flow within a single computation. The resultant intra-thrombus flow velocities are illustrated using thrombus cross-sectional slices taken along vessel axis (*sections labelled S*1 – *S*3) and perpendicular to the vessel axis (*labelled S*4 – *S*6) as shown in Figure 2, panel a. Intra-thrombus velocities for the heterogeneous thrombus microstructure considered here, are shown in panel d. for sections *S*1 – *S*3 and times *T*1 – *T*4; and in panel e. for sections *S*4 – *S*6 and times *T*1 – *T*4 respectively. Computed systolic shear strain rate at the vessel wall ranged between 0.10 sec^−1^ and 4.4 × 10^3^ sec^−1^. Maximum shear was located along the wall lateral to the thrombus, while the minimum was on vessel wall immediately distal to thrombus. Thrombus boundary shear strain rates ranged between 3.3 sec^−1^ and 5.5×10^3^ sec^−1^ (see *also supplementary material*).

**Figure 2:**
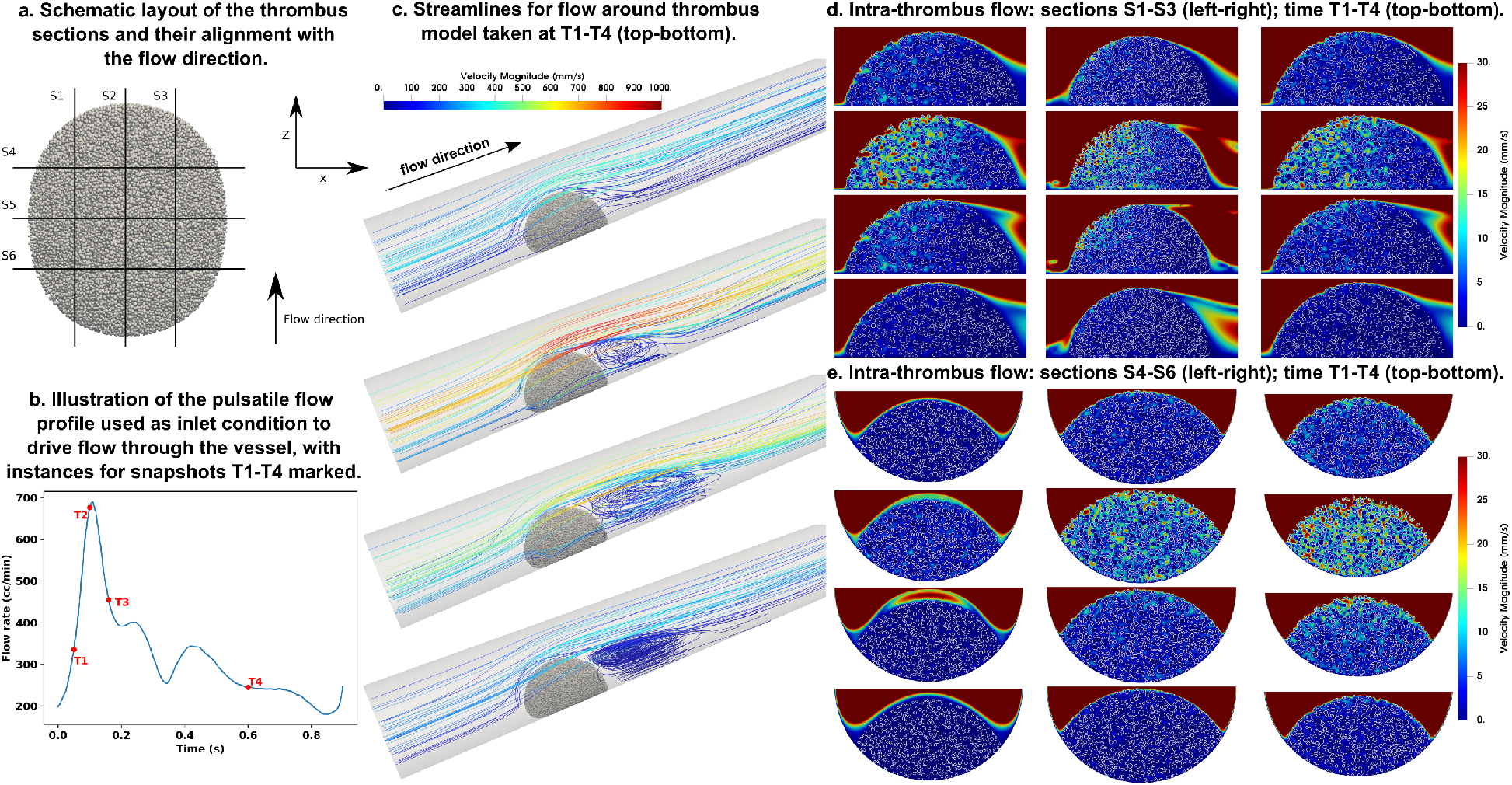
Illustration of computed unsteady flow within and around the thrombus. Four instances T1 – T4, marked on panel b., are chosen for visualization; and 9 sections S1 – S9 marked on panel a., are chosen for probing the thrombus interior flow field. Resulting unsteady streamline patterns are presented in panel c. Intra-thrombus velocities for lateral sections S1 – S3 are presented in panel d., and for axial sections S4 – S6 are presented in panel e.

Intra-thrombus velocities are observed to be ≥ 2 orders of magnitude smaller than extra-thrombus flow - consistent with previously demonstrated studies on microscale platelet plugs^[46,30]^ and two-dimensional macroscale models^[33,45]^. Specifically, we observe a permeation region along the boundary *shell* of the thrombus, which draws flow into the thrombus interior through pore network pathways. These permeation pathways are pronounced along routes linking to the upstream or proximal face of the thrombus, while distal regions have substantially slower flows. Beyond this permeation shell, the interior *core* region of the thrombus remains a region of arrested flow, with significantly low velocities, even during systole when most permeation flow occurs. The *core-shell* architecture is a common descriptor of thrombus microstructure originally derived from platelet packing phenomena in pathological clotting scenarios^[46,6]^. Our data on flow within a large artery thrombus hint that the level of hydraulic resistance offered to flow at the macroscopic scales (*beyond individual platelet scales*) also mimics this core-shell pattern. We note that resulting intrathrombus flow velocities for the macroscale thrombus is significantly higher compared to those reported in microscale thrombi, owing to overall faster external flow and regions of larger pore-sizes, especially in the shell. We remark that in prior work^[45]^ we demonstrated that flow leakage through thrombosed vessel wall substantially alters intra-thrombus flow characteristics - an additional aspect that we have not included in our three-dimensional model to keep within the scope of our study.

### 3.2 Drivers of three-dimensional flow mediated transport

Our treatment of the thrombus boundary as a heterogeneous permeable interface enables computing a normal thrombus boundary pressure gradient (TBPG) into the domain; which is recognized as a key factor for flow permeation and transport into the thrombus interior. In addition, quantification of forward-time FTLE fields enables assessment of local flow structures which form pockets or barriers organizing advective mass transport in the flow^[42,19]^. These quantifiers are employed to elucidate the drivers of flow-mediated transport in the extra-thrombus environment. Specifically, in Figure 3, spatial distribution of computed TBPG is presented along the thrombus boundary surface for instances *T*1 – *T*4 in the cardiac cycle. The gradients are viewed from the side facing the flow (top); the top (*middle*); and the distal side of the thrombus (*bottom*). The data clearly demonstrate that TBPG is non-uniform along the boundary surface, with peak TBPG occurring on the proximal side of the boundary at an angle from the base of the thrombus. This angle is governed, as viewed from Figure 2, by the unsteady flow structures formed due to thrombus-hemodynamics interactions, and flow interactions with the vessel wall. Observed TBPG values are maximal around systolic phase of the cardiac cycle, with peak systolic gradients reaching > 1.00 *mmHg/mm* while peak diastolic gradients being ≈ 0.10 *mmHg/mm*. FTLE fields for the corresponding time instances *T*1 – *T*4 are illustrated in Figure 4. The peak and trough of the spatially varying FTLE fields are volume rendered to illustrate the threedimensional unsteady coherent structures forming around the thrombus due to thrombus-hemodynamics interactions (*top illustration in each panel*). A coherent envelope is observed around the thrombus which strengthens during systole, and decays during diastole. The envelope has a prominent component proximal to the thrombus at all times; in addition to the time-varying component in the distal region as illustrated by sectional views of the FTLE fields taken along the vessel axis (*bottom illustration on each panel*). We have overlaid the TBPG along thrombus manifold surface with FTLE sectional views - illustrating that the region where proximal FTLE ridges couple to the thrombus boundary correlate with regions of high TBPG.

**Figure 3:**
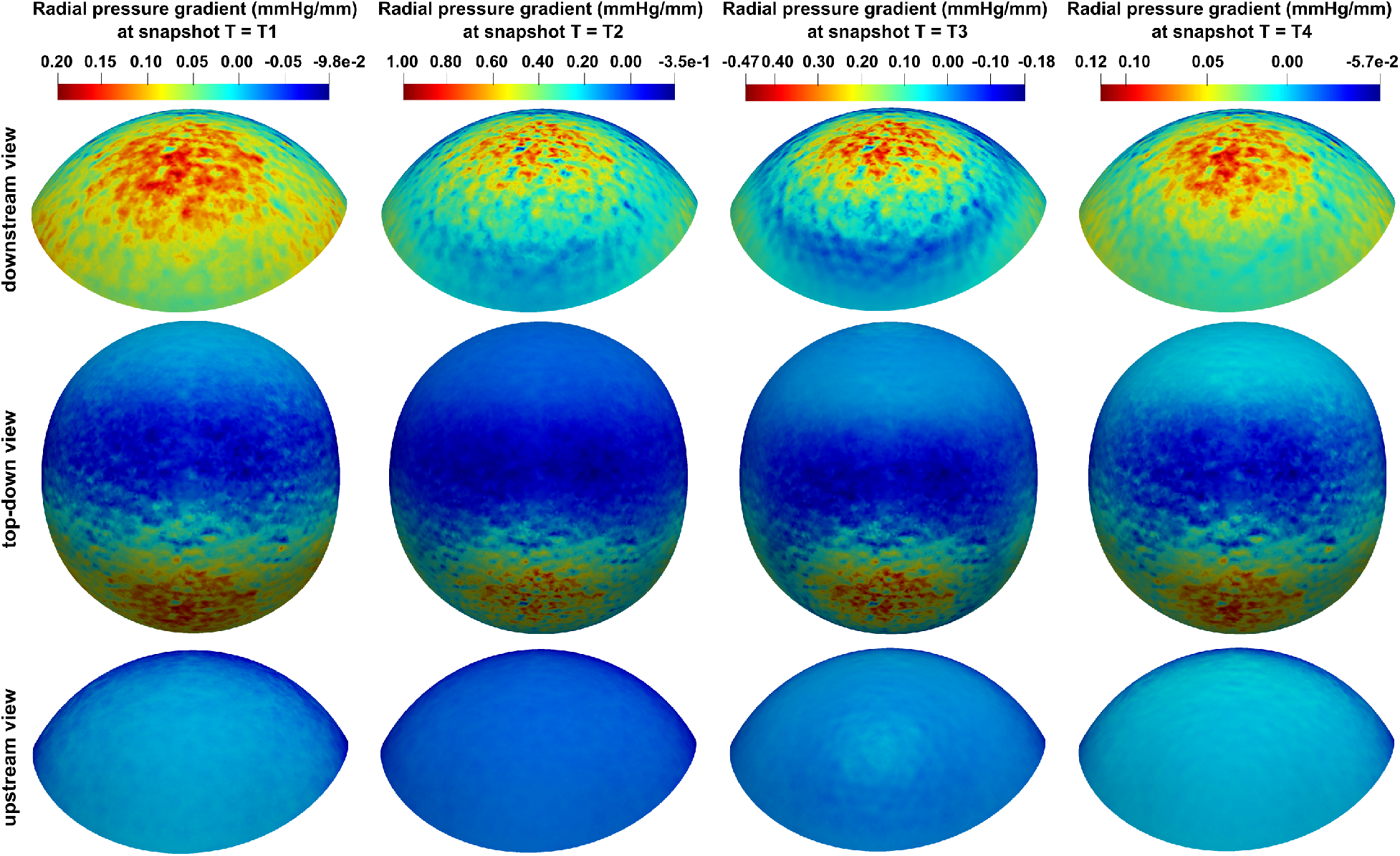
Computed values of thrombus boundary pressure gradient (TBPG) viewed at instances T1 – T4 from the side proximal to the flow (top); from the top (middle); and from the side distal to the flow (bottom).

### 3.3 Transport of species mixed in flow from upstream

To characterize how blood flow transports biochemical species towards the thrombus and in the etxra-thrombus environment, we simulated the transport of a non-reactive biochemical species with diffusivity *D* = 5.0 × 10^−5^ *mm*^2^/*sec* with a fixed concentration *c* = 2.0 *μ*Molar specified as boundary condition at vessel inlet. For this diffusivity and the flow regime in the vessel around the thrombus, we obtain an advection dominant transport (*high Peclet number*) regime. The advection dominant process is expected, therefore, to follow the coherent structure identified by the FTLE field, which is what we observe from the spatial variation of the concentration data illustrated in Figure 5 (*time T1-T4 presented top-bottom*). The figure presents a lateral view of the volume rendered concentration field (*in yellow*), superimposed with the coherent structures depicted in Figure 4. We observe that the concentration field obeys the barrier envelope formed by the high FTLE value - advective species transport does not cross the coherent envelope around the thrombus. While the presence of the FTLE envelope prevents material from the bulk of the flow reaching the thrombus boundary for most of the thrombus domain, the portion of the envelope proximal to the thrombus as discussed before does bring in species released from the inlet towards the thrombus. This species concentration reaches the proximal side of the thrombus boundary, and permeates into the thrombus interior driven by the high TBPG associated with this region (see Figure 3). To characterize this intra-thrombus permeation, we projected the concentration values on the surface of concentric fixed-radius shells starting from the outer boundary of the thrombus. The resulting concentration fields are illustrated in Figure 6. The shells illustrated here are taken from the outer boundary (*r* = 3.0 *mm*) until 0.20 *mm* inside the thrombus, in increments of 0.05 *mm*. The first row of the shell depictions in Figure 6 clearly show the microstructural heterogeneity by superposing the particles (*seen here for the boundary*) onto the shell, and establishing that the concentration permeation occurs along the pore network pathways around the discrete particles into the thrombus interior. Variation in time *T*1 – *T*4 is shown from left to right, and variation in shell depth is shown across rows 2-6. In accordance with the points mentioned above, a correlation of the pressure gradient and the concentration on the outer surface is observed, as well as the heterogeneity of the permeation transport into the thrombus interstices is evident.

**Figure 4:**
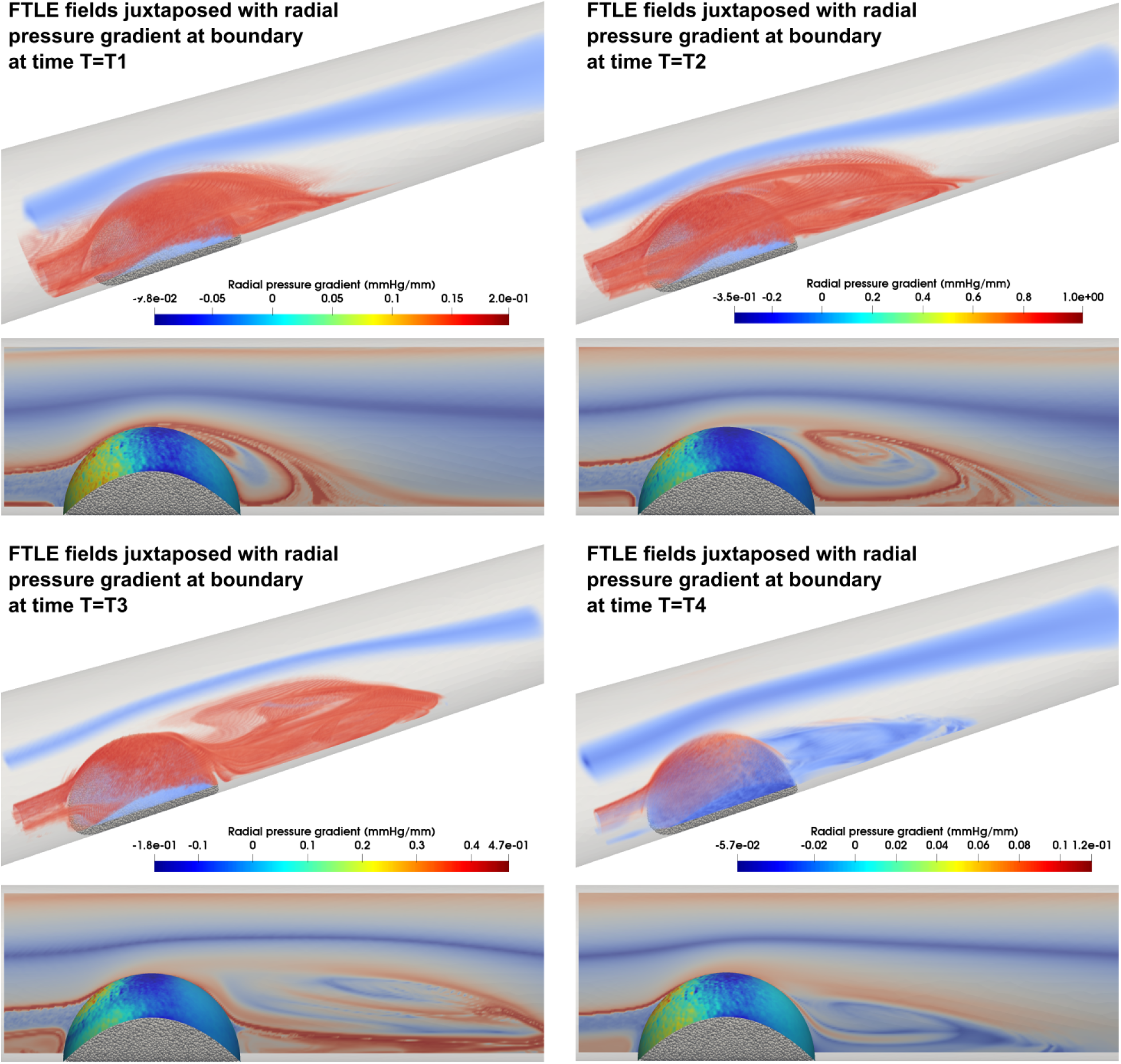
Illustration of FTLE fields generated by unsteady thrombus-hemodynamics interactions for times T1 – T4. Top view in each panel shows the coherent structure that envelopes the thrombus and evolves over time. Bottom view in each panel depicts how FTLE correlates to TBPG peak regions. FTLE peaks and troughs are denoted in red and yellow respectively in the top view.

**Figure 5:**
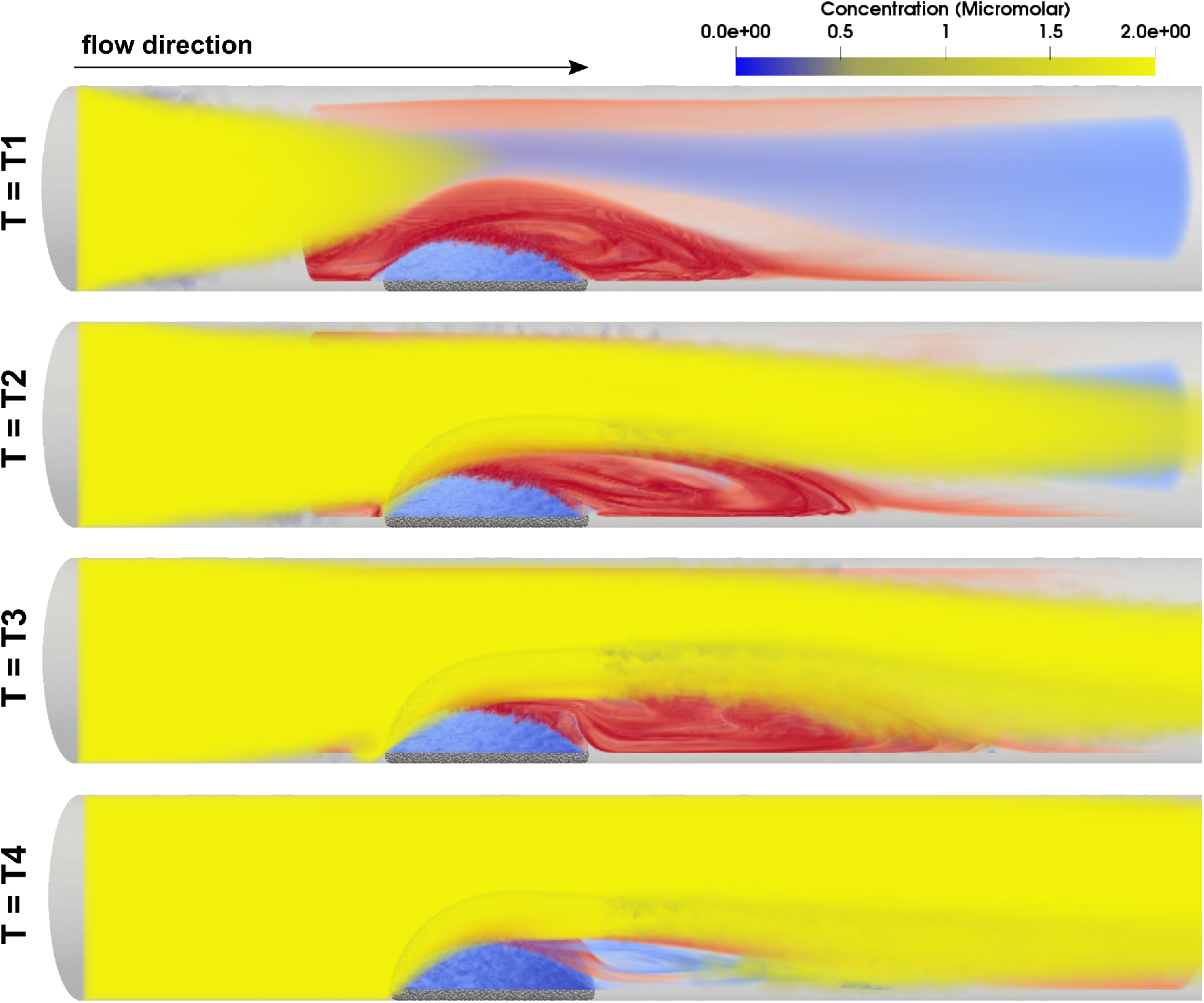
Volume rendered time-evolving concentration field of species with D = 5 × 10^−5^ mm^2^/sec (yellow), superposed with the coherent structures around the thrombus identified by FTLE peaks/troughs (red/blue). A fixed concentration 2.0 μ Molar is specified herer at vessel boundary.

**Figure 6:**
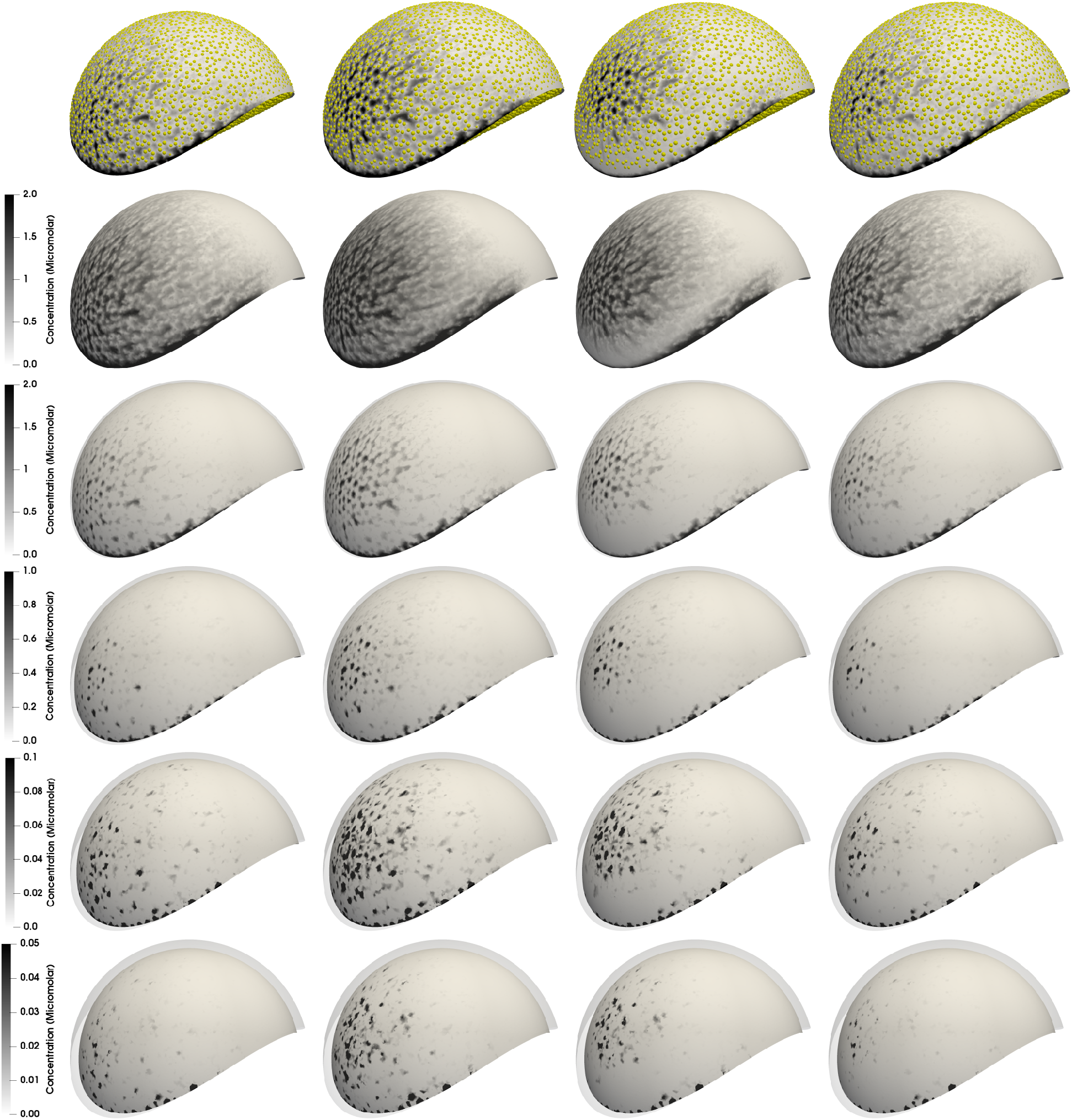
Heterogeneous permeation of species with D = 5 × 10^−5^ mm^2^/sec visualized for radially concentric shells taken depthwise into the thrombus starting from the boundary (row 2) upto 0.2 mm into the thrombus (row 6). Top row shows a visualization of the heterogeneity in microstructure that manifests in the resulting spatial concentration distribution.

### 3.4 Intra-thrombus transport and species concentration

Following flow-mediated permeation into the thrombus, subsequent intra-thrombus transport is primarily diffusion dominant process, indicated by slow flow velocities as seen in Figure 2 and small interstitial spaces (*together leading to low Peclet number*). The long time-scale behavior of this diffusive transport, and intra-thrombus concentration variation, are studied using a time-averaged fictitious domain AD simulation. Specifically, the thrombus domain mesh is geometrically extracted, and flow velocities on the mesh are averaged across the final simulated cardiac cycle. Concentration at the thrombus boundary brought in from the vessel inlet (as *shown in Figure 6*) is also cycle-averaged, and assigned as a Dirichlet boundary condition at the boundary nodes of the mesh for a steady-state AD simulation. A series of time-averaged computations were run for species diffusivity values ranging from *D* = 5 × 10^−6^ *mm*^2^/*sec* to *D* = 5 × 10^−1^ *mm*^2^/*sec*, varied by factors of 10. The simulated concentration values within the heterogeneous thrombus domain were then averaged along circumference of concentric spherical shells, starting from the thrombus boundary and progressing in increasing order of depth into the thrombus. The resulting shell-averaged concentration variation curves are presented in Figure 7. The y-axis is normalized against boundary concentration, providing a non-dimensional characterization of intra-thrombus transport with varying species diffusivity. For values lower than *D* = 5 × 10^−6^ *mm*^2^/*sec* the concentration gradients did not change appreciably owing to the associated Peclet number becoming high enough to shift the dominant transport modality to advection. We note that owing to the heterogeneous microstructure model, and associated heterogeneities in pore-size and flow velocities (*eg. core-shell distinction*) quoting a single characteristic length and velocity scale - and consequently, a single Peclet number - for the intra-thrombus transport is difficult. Hence, in Figure 7, we simply characterize the data in terms of values of *D* directly.

**Figure 7:**
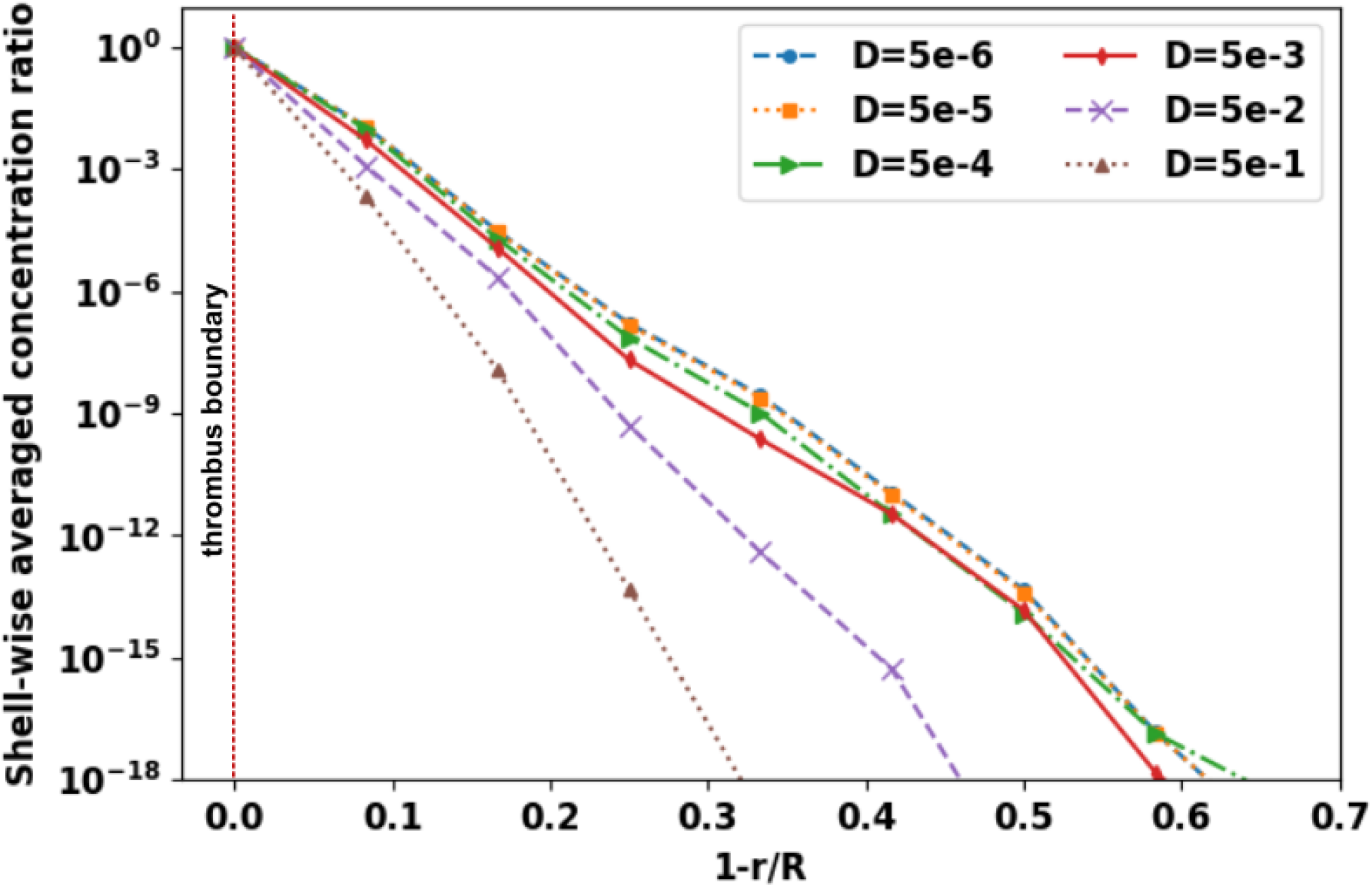
Concentration values averaged on radially concentric shells taken depthwise into the thrombus starting from the outer boundary; for varying diffusivity.

### 3.5 Transport of species released from thrombus domain

Transport of species emanating from the thrombus boundary was studied by releasing a fixed concentration of 2.0 *μ*Molar from discrete particles within a shell of 0.20 *mm*. The resultant concentration fields for species diffusivity value of *D* = 5 × 10^−5^ *mm*^2^/*sec* are illustrated in Figure 8 in panel b. (top view of the thrombus neighborhood at section midway through the depth of the thrombus). The corresponding FTLE fields, with peaks and troughs, are shown on panel a. To further illustrate the transport behavior, a second case with diffusivity *D* = 5 × 10^−2^ *mm*^2^/*sec* was also simulated, and the resultant concentration distributions are shown on panel c. We observe that concentration remains prominently trapped within the coherent structure formed adjacent to the thrombus (panel a) at all instances. Higher extent of diffusion enables more concentration to exit these structures into the bulk flow (panel c), which leads to reduced sharpness of concentration gradients, and reduced amount of concentration pooling. However, the bulk of the concentration still remains adjacent to the thrombus within the coherent structure formed distal to the thrombus domain. These aspects indicate that for physiologically relevant diffusivity ranges, material released from the thrombus boundary will tend to follow near-thrombus coherent structures originating from the thrombus-hemodynamics interactions.

**Figure 8:**
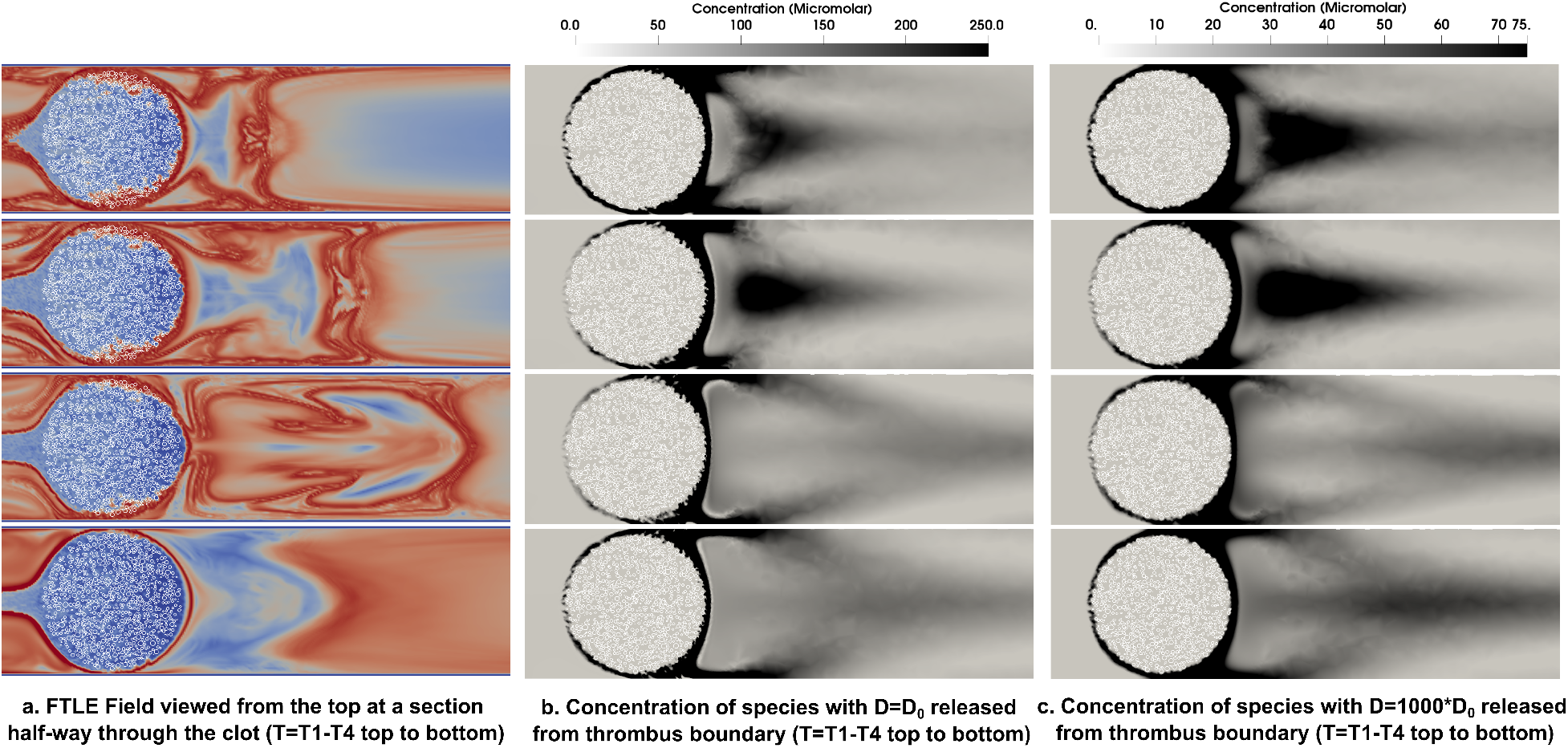
Concentration fields for species with diffusivity D = 5 × 10^−5^ mm^2^/sec (panel b.) and D = 5 × 10^−2^ mm^2^/sec (panel c.) compared with corresponding FTLE peaks/troughs (panel a.) for cases where 2.0 μMolar was released at the thrombus boundary and run for 40 cardiac cycles. Section views taken at half-way through depth of thrombus. FTLE peaks and troughs are denoted in red and yellow respectively in panel a.

## 4 Discussion

Results from our numerical investigations elucidate the fundamental factors determining the state of flow-mediated transport for large artery thrombi. Specifically, outside the thrombus in the vessel, advective transport brings species into the thrombus neighborhood, where thrombus-hemodynamics interactions lead to formation of dynamic coherent structures that organize the advective phenomena. Permeation flow, driven by a pressure gradient originating also from unsteady flow interactions, then leads to species transport from the thrombus neighborhood into the thrombus interior. Once permeated into the interior, advective processes gradually get dominated by diffusive processes until into the thrombus core where flow is arrested, and transport is diffusion dominant. Permeation processes at the boundary therefore play a central role, and our methodology enables efficient modeling of realistic heterogeneous thrombus boundary for detailed characterization of permeation. We note that microstructural heterogeneities also determine the intra-thrombus transport pathways post-permeation, and ultimately govern the balance between advection-diffusion transport in thrombus interior. We have demonstrated here that our microstructure aware approach enables detailed three-dimensional investigations of these aspects, which are otherwise challenging to conduct.

Efficient resolution of thrombus-hemodynamics interactions is essential to identify coherent structures organizing advection around the thrombus. We demonstrated here that the presence of these coherent structures may mean that: (a) all material from the bulk flow does not find its way to the thrombus boundary; and likewise (b) all material released from thrombus boundary may not find its way into the bulk flow. Thrombus deformation in response to hemodynamic loading can further influence these local coherent structures, an aspect that is substantially complex to address within the scope of this study. However, thrombus formation is commonly followed by a phase of retraction and thrombus consolidation driven by platelet contractile forces^[7,21,8]^, which renders stability and stiffness and reduces the extent of thrombus deformation. The stable, compact thrombus aggregate motivates our no-deformation assumption for this study. To the best of our knowledge, this is one of the early comprehensive attempts at characterizing the fully three-dimensional unsteady coherent structures around a heterogeneous thrombus aggregate, and establish their role in thrombotic phenomena.

Consideration of physiologically realistic diffusivity regimes of 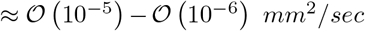 also reveals an interesting aspect. Estimating diffusion time scale as *T_D_* ≈ *x*^2^/4*D* for a characteristic distance *x*, diffusion dominant transport across 1.0 *mm* macroscopic length is expected to range anywhere between 7 hours to 2.5 days. One cardiac cycle is ≈ 1.0 *sec*, and commonly simulated using numerical time-steps ≈ 1/1000’th of a cycle. Advection, driven by the bulk flow, also proceeds on similar time-scales. This establishes the spatiotemporally multiscale nature of transport in and around a macroscopic large artery thrombus. Having illustrated this, we demonstrated how to address the separation of time-scales numerically by conducting an unsteady shorter time-scale advection resolved simulation, and feeding the resulting thrombus boundary concentration into a longer-scale time-averaged diffusion resolved simulation.

The microstructure aware approach and associated findings presented here have several physiological implications. One critical implication pertains to mechanism and efficacy of thrombolytic drug transport. Thrombolysis is essentially a spatially non-uniform, heterogeneous process. The amount and spatial extent of drug(tPA) transport into the thrombus interior is a key determinant of treatment success. Our findings clearly indicate that TBPG driven permeation into the shell will initiate the process, and drug concentration will drop off quickly across the depth of the thrombus. However, as the fibrinolytic reactions progress, more of the shell will lyse, exposing more of the interior through which drug can permeate further inwards. Our approach, therefore, enables studying the flow-mediated mechanisms of thrombolysis, and provides an effective numerical avenue to develop detailed *in silico* models of thrombolysis. In addition, our findings also have implications for the transport of coagulation agonists like ADP and Thromboxane released from the thrombus boundary. While these agonists play a role in further activating and recruiting platelets thereby enabling disease progression, our findings hint that depending upon the thrombus-hemodynamics interactions, their transport in the bulk flow may be restricted significantly by the local coherent structures around the thrombus.

While the geometric simplifications here enabled convenience of parametric simulations, extension to physiologically realistic vascular anatomy and thrombus geometries are the necessary next steps. Realistic vessel geometries have curvature induced vortical flows which will promote mixing and, together with interactions with realistic thrombus shapes, determine more realistic FTLE and TBPG values. Furthermore, elevated shear rates in the thrombus neighborhood may lead to local fluctuations in blood rheology. Accounting for such local effects requires complex, non-trivial extension of the fictitious domain finite element framework^[33,45]^ to incorporate non-Newtonian rheology, which was not the scope of this present study. Additionally, while we considered passive species here, most relevant biochemical species will have associated roles in biochemical reactions which need to be accounted for to advance these models further. Finally, twoway coupled interactions between unsteady hemodynamics and thrombus deformation mechanics requires considerable computational challenges, which necessitate separate discussion of its own, and have not been addressed here to keep within the scope of the study. Nevertheless, the numerical machinery developed here, and the underlying mechanistic aspects of flow-mediated transport retain their significance for large artery thrombi.

**5 Concluding Remarks**

We presented a custom preconditioned fictitious domain model for thrombus-hemodynamics interactions coupled with a fictitious domain advection-diffusion model for biochemical species transport and a discrete particle representation of heterogeneous thrombus microstructure. The resultant approach enabled microstructure aware investigations on flow and transport within and around large arterial thrombi; which we demonstrated using a geometrically simplified model system of a symmetric hemispherical thrombus embedded in a cylindrical vessel. Simulations on the model system illustrated the fundamental processes underlying flow-mediated transport around the thrombus, permeation into the thrombus, and subsequent intra-thrombus transport.

## 7 Acknowledgements

This work utilized resources from the University of Colorado Boulder Research Computing Group, which is supported by the National Science Foundation (awards ACI-1532235 and ACI-1532236), the University of Colorado Boulder, and Colorado State University. The Authors also gratefully acknowledge guidance, support, and the many valuable discussions with Prof. Scott L. Diamond, Department of Chemical and Biomolecular Engineering, University of Pennsylvania; and Prof. Shawn C. Shadden, Department of Mechanical Engineering, University of California, Berkeley. These fruitful discussions strongly benefited the study design and interpretation of results. CT implemented the computer algorithms, performed the flow simulations, and data analysis. DM developed the numerical methods and algorithms, designed the study, and wrote the manuscript. All authors reviewed the manuscript and agreed to the final version.

## Supplementary information

Here we include additional relevant details regarding our computational approach described in this article. Details regarding the PCD-FD approach, and the mathematical details regarding the kernels used for fictitious domain AD formulations are included. Additionally, for the sake of rigor and completion, we have included sample numerical validation cases for our fictitious domain AD implementation. The corresponding validation cases for the flow velocity formulation have already been provided in prior work. Lastly, we include a few pieces of supplementary data to support the content presented in the main manuscript.

### Details of PCD-FD block matrix system contributions

To illustrate the role of fictitious domain contribution to the block matrix system, the variational problem must first be linearized via Newton’s iteration. Since the boundary integral term and stabilization term does not affect the fictitious domain contribution, they are omitted in this analysis. The variational problem with fictitious domain contribution then becomes

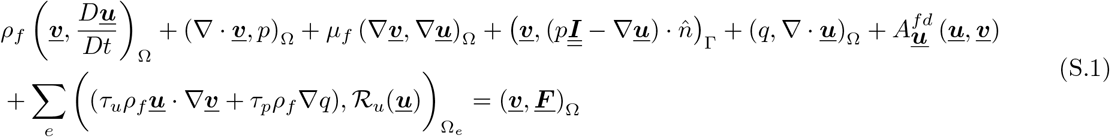

Using the *θ* time integration scheme at time step *n*, the system becomes

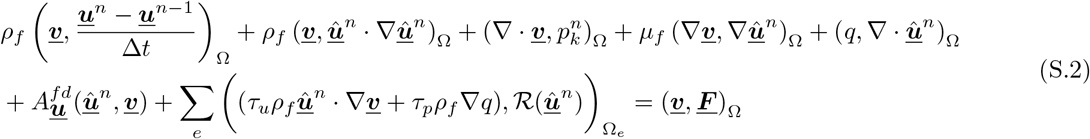

where 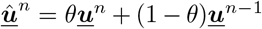 and 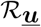 is the residue of the momentum equation. Using Newton’s iteration step *k*, the solution 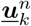 is updated as

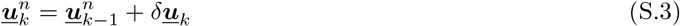

using the definition of 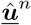 we get the definition of 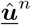 at Newton’s iteration step *k* as

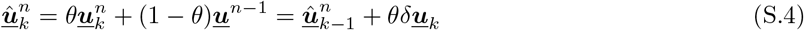

By applying 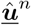 and only keeping only the first order terms, the system becomes

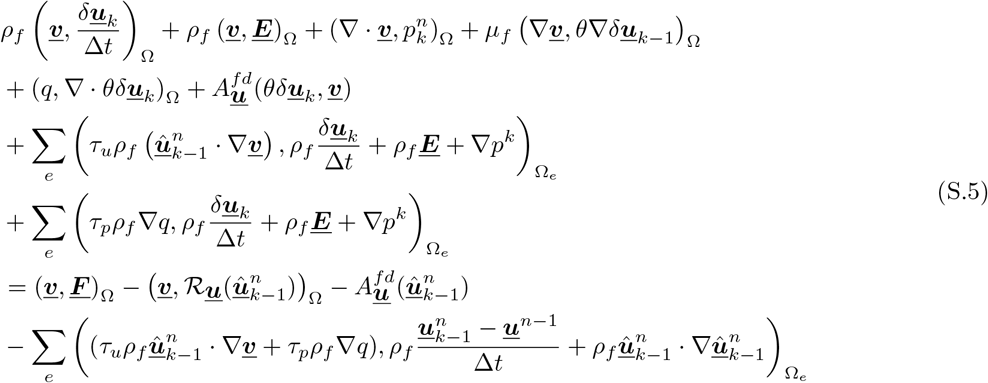

where

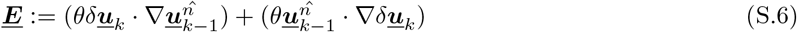

Note that the stabilization parameter *τ_u_* and the Newton iteration increment 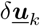 are small therefore the terms with 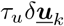 are considered small compared to other terms and drop out. 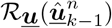 is the residue of the variational problem evaluated as the previous Newton iteration solution 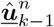. Note that since the fictitious domain contribution is linear in ***<u>u</u>***, no additional terms arise from linearization. The left hand side of equation (S.5) can be written in a block matrix form as:

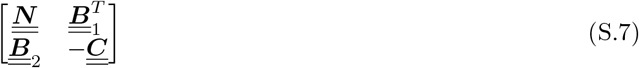

where

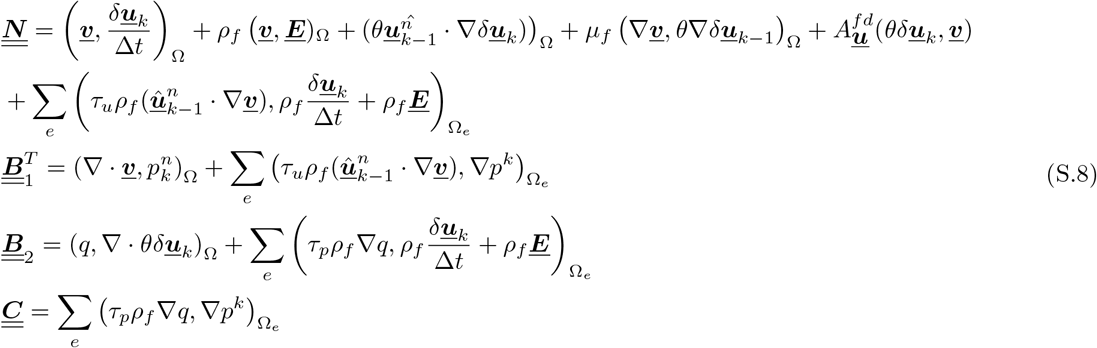

In the PCD preconditioner formulation, the fictitious domain contribution 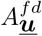 is added in the construction of the projected convection-diffusion operator 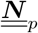 onto the pressure function space.

### Discrete particle kernel selection

For the kernel function 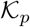 as shown in Equation 10, we employed a radially truncated Gaussian kernel function - defined on the particle domain 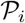 as:

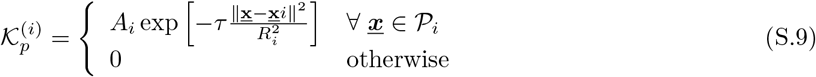

where ***<u>x</u>_i_*** is particle’s center, *R_i_*, is the particle’s radius, *τ* controls the width of the kernel, and *A_i_* is chosen such that the integral over the particle’s domain is normalized.

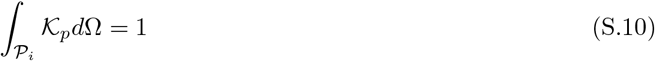

For a spherical particle, the normalization factor can be analytically obtained as:

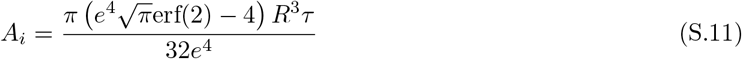

Implementation of this kernel on a collection of particles as utilized in our fictitious domain AD formulation is shown in Figure S1.

**Figure S1:**
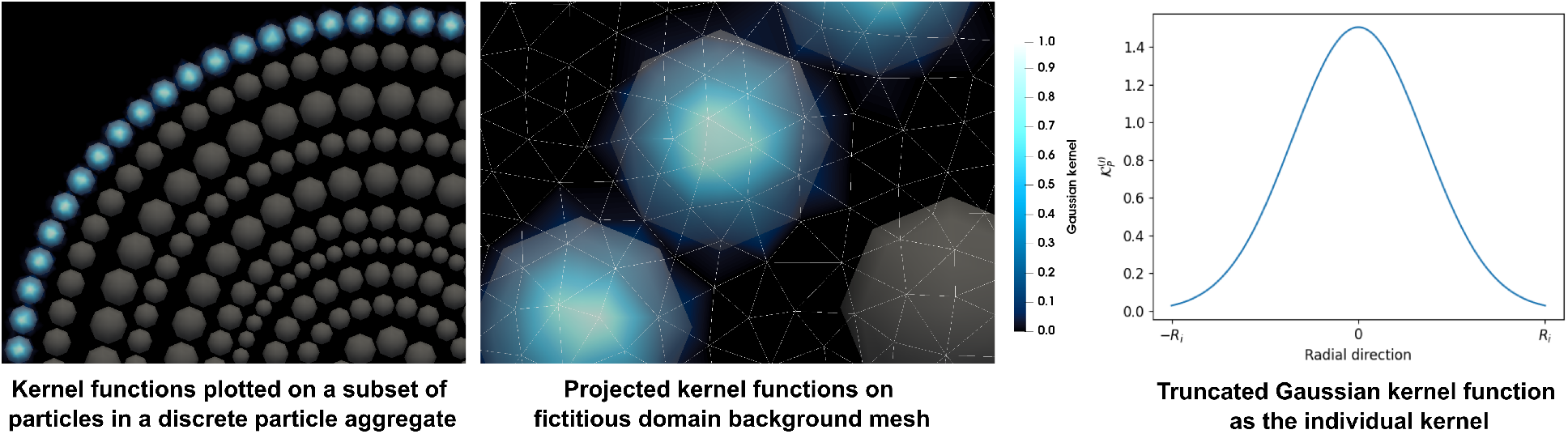
Illustration of the discrete particle kernel implementation for fictitious domain AD formulation.

### Validation of fictitious domain approach for transport problems

We conducted validation studies for the fictitious domain approach for transport problems on a square domain with a monodisperse particle packing arrangement obtained by a random sequential addition process. We compared our fictitious domain approach against the case where the particle’s boundary is meshed explicitly, and concentration boundary values are set explicitly using a Dirichlet boundary condition. For a sample validation case, we consider a diffusion problem on the domain and a Dirichlet boundary condition is applied at the bottom of the domain (*c* = 2.0*μ*Molar) and a homogeneous boundary condition is applied on the particle’s boundary and the top boundary of the domain. The diffusion coefficient is *D* = 5 × 10^−5^ *mm*^2^/*s* which match the physiological range considered in this work. The results are shown in Figure S2 and clearly establishes that the fictitious domain approach is in good agreement with the case of explicitly prescribed boundary condition on the particle’s boundary.

**Figure S2:**
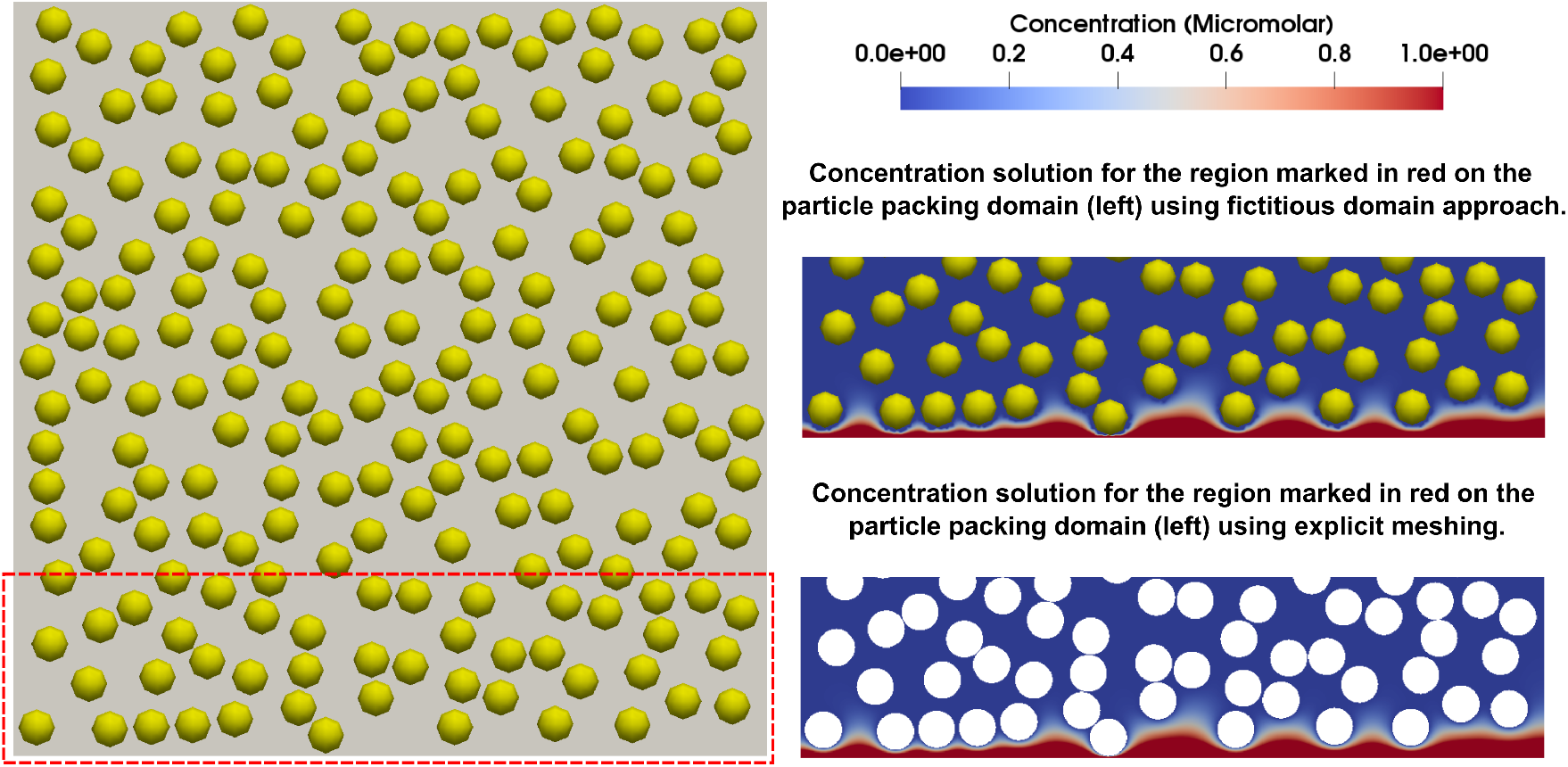
Validation of fictitious domain approach to transport problems - comparing boundary condition on the particles imposed using fictitious domain approach with those imposed explicitly on the particles.

### Supplementary data: Statistics of discrete particle aggregate model

The discrete element representation of the heterogeneous thrombus microstructure comprises spherical polydisperse particles generated as discussed in Section 2.4. The histogram depicting size distribution of the polydisperse discrete particles is provided in Figure S3 below. The mean particle/element diameter is 98 microns, and the mode (that is, most probable) diameter is 77 microns. Assuming simplified Kozeny-Carman relation, we can obtain a baseline permeability estimate for the polydisperse aggregate (which is only to serve as a reference, not as an exact estimate of thrombus hydraulic permeability):

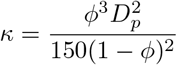

where *ϕ* is average porosity (0.68 for our case) and *D_p_* is average diameter. With these values we obtain a reference permeability of approx. 197 micron^2^.

**Figure S3:**
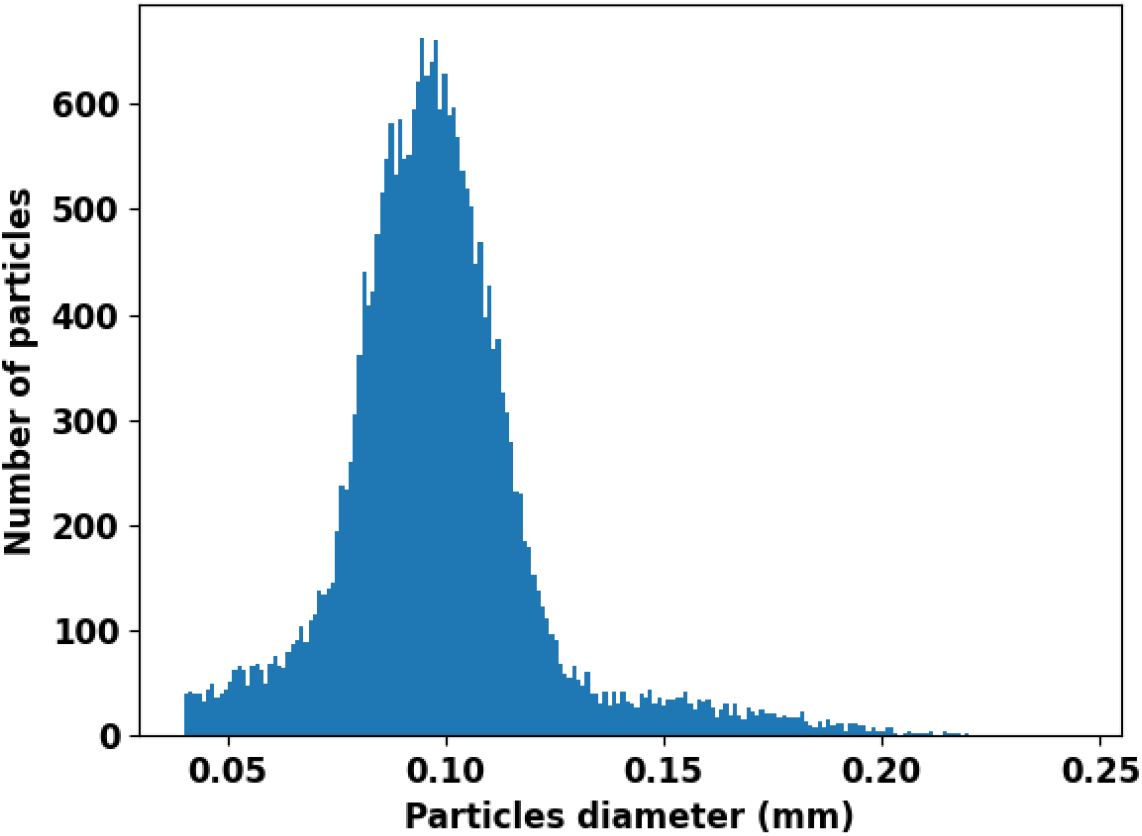
Histogram of discrete element/particle sizes used for heterogeneous thrombus microstructure reconstruction

### Supplementary data: Shear on thrombus boundary and vessel wall

Computed wall shear strain rates for locations along the vessel wall have been compiled in Figure S4 below. In panel a and b, we depict the systolic wall shear strain rates, while in panel c. we depict a reduced strain-rate scale to identify the regions of low wall shear strain rate distal to the thrombus. Similarly, in Figure S5 we depict the systolic wall shear strain rates for the thrombus boundary (visualized on a hemispherical shell manifold outlining the thrombus boundary). Front (or proximal) view along flow direction, top-down view, and rear (or distal) view against flow direction has been depicted in panels a., b., and c. respectively.

**Figure S4:**
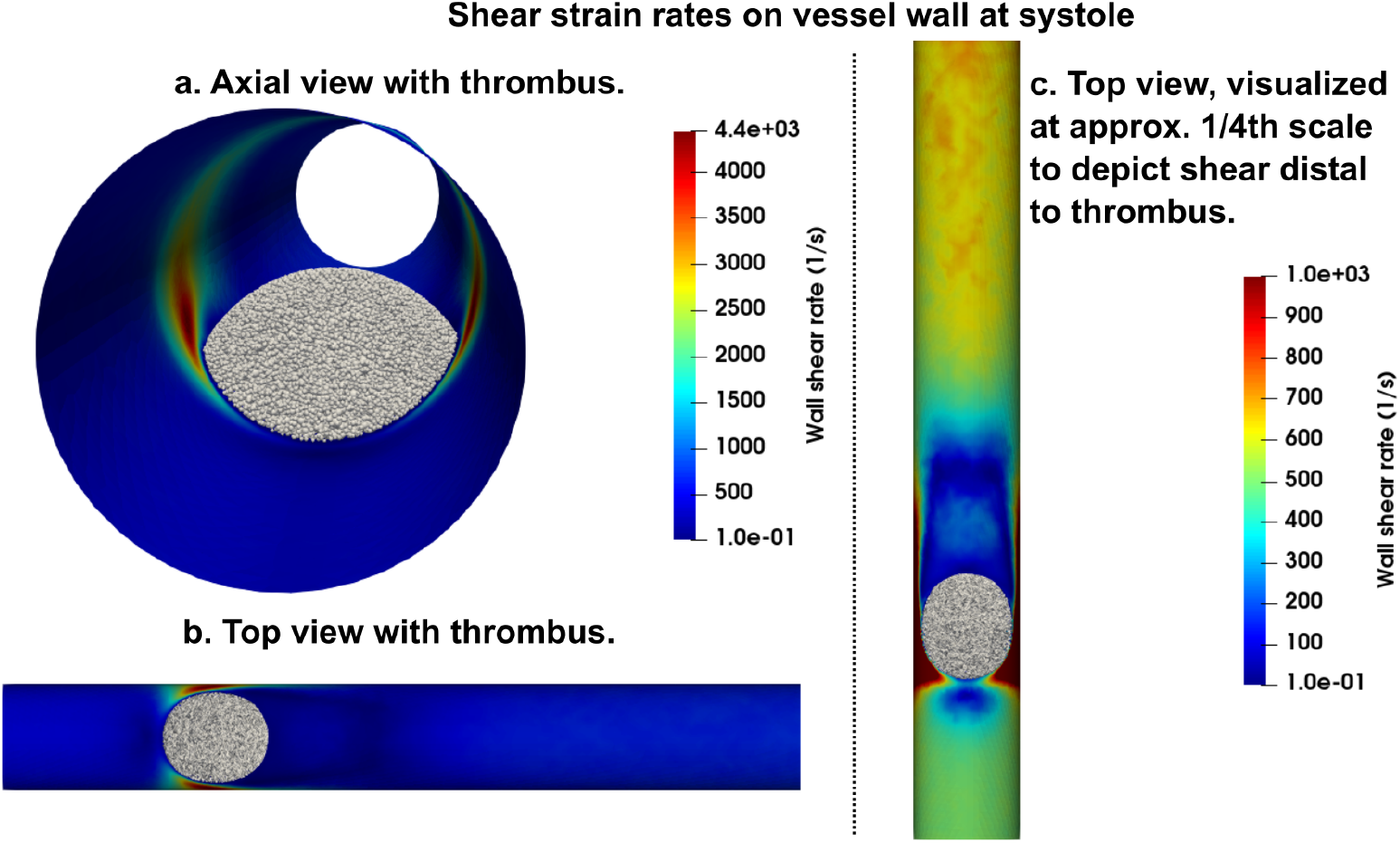
Wall shear strain rate on the vessel wall at peak systole, visualized with the thrombus in place.

**Figure S5:**
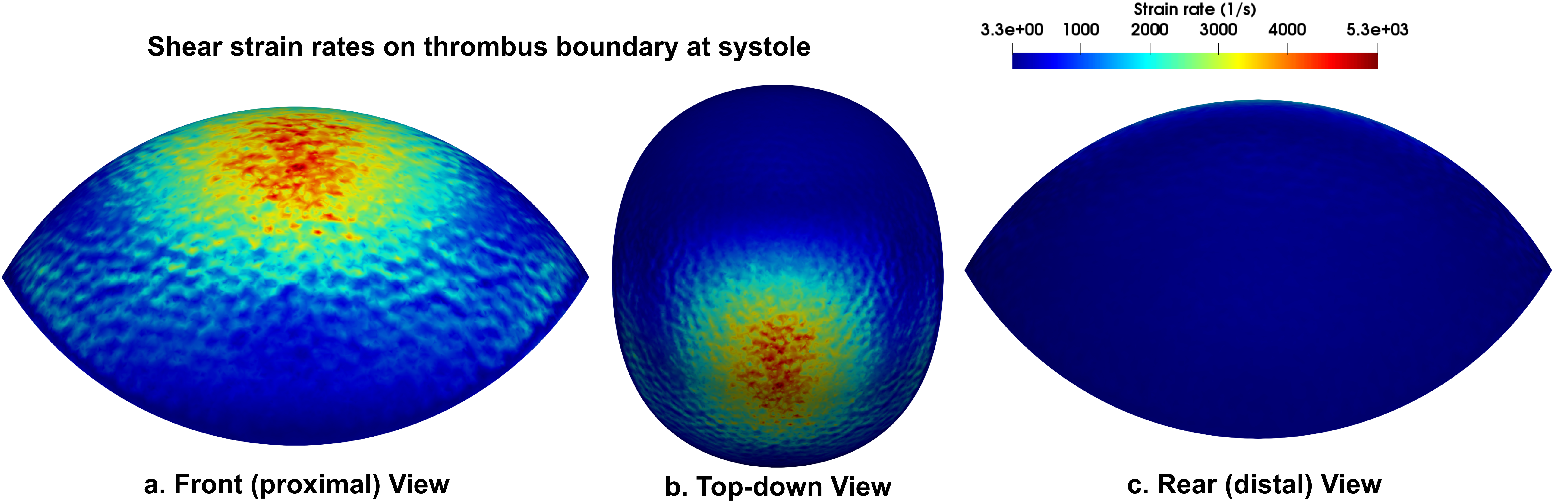
Wall shear strain rate on the thrombus boundary at peak systole.

